# Associations Between Prenatal Vitamin D and Placental Gene Expression

**DOI:** 10.1101/2024.05.10.593571

**Authors:** Mariana Parenti, Melissa M. Melough, Samantha Lapehn, James MacDonald, Theo Bammler, Evan J. Firsick, Hyo Young Choi, Karen J. Derefinko, Daniel A. Enquobahrie, Kecia N. Carroll, Kaja Z. LeWinn, Nicole R. Bush, Qi Zhao, Sheela Sathyanarayana, Alison G. Paquette

## Abstract

**Background:** Vitamin D is a hormone regulating gene transcription. Prenatal vitamin D has been linked to immune and vascular function in the placenta, a key organ of pregnancy. To date, studies of vitamin D and placental gene expression have focused on a limited number of candidate genes. Transcriptome-wide RNA sequencing can provide a more complete representation of the placental effects of vitamin D.

**Objective:** We investigated the association between prenatal vitamin D levels and placental gene expression in a large, prospective pregnancy cohort.

**Methods:** Participants were recruited in Shelby County, Tennessee in the Conditions Affecting Neurocognitive Development and Learning in Early childhood (CANDLE) study. Vitamin D level (plasma total 25-hydroxyvitatmin D, [25(OH)D]) was measured at mid-pregnancy (16-28 weeks’ gestation) and delivery. Placenta samples were collected at birth. RNA was isolated and sequenced. We identified differentially expressed genes (DEGs) using adjusted linear regression models. We also conducted weighted gene co-expression network analysis (WGCNA).

**Results:** The median 25(OH)D of participants was 21.8 ng/mL at mid-pregnancy (*N*=774, IQR: 15.4-26.5 ng/mL) and 23.6 ng/mL at delivery (*N*=753, IQR: 16.8-29.1 ng/mL). Placental expression of 25 DEGs was associated with 25(OH)D at mid-pregnancy, but no DEG was associated with 25(OH)D at delivery. DEGs were related to energy metabolism, cytoskeletal function, and RNA transcription. Using WGCNA, we identified 2 gene modules whose expression was associated with 25(OH)D at mid-pregnancy and 1 module associated with 25(OH)D at delivery. These modules were enriched for genes related to mitochondrial and cytoskeletal function, and were regulated by transcription factors including *ARNT2*, *BHLHE40*, *FOSL2*, *JUND*, and *NFKB1*.

**Conclusions:** Our results indicate that 25(OH)D during mid-pregnancy, but not at delivery, is associated with placental gene expression at birth. Future research is needed to investigate a potential role of vitamin D in programming placental mitochondrial metabolism, intracellular transport, and transcriptional regulation during pregnancy.

## 1. INTRODUCTION

During fetal development, vitamin D is involved in skeletal formation and growth, immune regulation, and placentation (1). Maternal vitamin D deficiency during pregnancy is associated with increased risk of adverse pregnancy and birth outcomes, including low birth weight, small for gestational age, preeclampsia, and gestational diabetes (2–5). Prenatal vitamin D may also program later-life health, including neurological and cardiometabolic health, through its role in endocrine function (6). Vitamin D deficiency and inadequacy during pregnancy and lactation is prevalent throughout the world, reaching approximately 33% in the United States (7,8).

Vitamin D is a preprohormone found in the diet. Humans can also synthesize it endogenously in their skin. Vitamin D is hydroxylated twice: first in the liver to the prohormone 25-hydroxyvitamin D [25(OH)D] and then in the kidney to the active hormone 1,25-dihydroxyvitamin D [1,25(OH)_2_D] by CYP27B1 (9). During pregnancy, the placenta is a key site of 1,25(OH)_2_D activation (10). The placenta expresses both vitamin D activating *CYP27B1* and the vitamin D-inactivating *CYP24A1*, and the vitamin D receptor (*VDR*) (10). This suggests that the placenta is capable of local vitamin D homeostasis and the vitamin D signaling might play an important role in the placenta (10). Vitamin D treatment promotes trophoblast invasion accompanied by increased matrix metalloprotease expression and secretion in primary human extravillous trophoblasts isolated from first trimester placentas and cell lines (11,12). Vitamin D also promotes amino acid transporter expression, which might link vitamin D status and fetal growth through nutrient transport via these receptors (13). Additionally, vitamin D deficiency activates the Hippo signaling pathway, which controls organ size during development, and was linked to fetal growth restriction in a rat model (14). Vitamin D treatment is also associated with reduced inflammatory signaling. Compared to untreated controls, 1,25(OH)_2_D treatment dose-dependently downregulated mRNA and protein expression of the inflammatory cytokines TNF-α and IL-6 in placentas from preeclamptic pregnancies (15). Similarly, over-representation analysis identified that the inflammation and immune regulation pathway was enriched for downregulated genes associated with 25(OH)D treatment in primary villous fragments from treated term human placenta compared to untreated controls (10). Accordingly, vitamin D could modulate pathways central to placental function as a conduit for fetal nutrition and a regulator of maternal-fetal immune interactions.

Candidate gene-based studies have revealed associations between placental gene expression and vitamin D supplementation or status, including positive associations with placental amino acid transporter gene expression (16), downregulation of an antiangiogenic factor associated with preeclampsia (17), and regulation of the inflammatory response after an immune challenge (18). In one recent transcriptome-wide study, short-term vitamin D treatment was associated with tissue remodeling and gene transcription in both transcriptomic and proteomic analysis in primary villous tissue isolated from human placentas (10). Additionally, vitamin D deficiency has been shown to elicit sex-specific effects on placental candidate gene expression in a mouse model, including *Vdr*, *Cyp24a1*, and *Cyp27b1* (18). Furthermore, testosterone might influence placental vitamin D metabolism, as it has been shown to downregulate gene expression of vitamin D-activating *CYP27B1* and upregulate expression of vitamin D-inactivating *CYP24A1* (19). However, the relationship between prenatal vitamin D levels and transcriptome-wide placental gene expression remains to be studied in an epidemiological context. Thus, we aimed to investigate the association between maternal vitamin D status during pregnancy and human placental gene expression, as well as the role of fetal sex as an effect modifier in a large, diverse prospective birth cohort.

## 2. METHODS

### 2.1. Study participants and data collection

This analysis was conducted using samples collected as part of the CANDLE (Conditions Affecting Neurocognitive Development and Learning in Early childhood) Study. This prospective birth cohort conducted in Shelby County, Tennessee has been described in detail elsewhere (20). Between December 2006 and July 2011, 1,503 pregnant participants were recruited during their second trimester and were considered eligible if they were between 16 and 28 weeks of gestation, had an uncomplicated singleton pregnancy, and planned to give birth at one of five participating Shelby County health care centers. Participants were included in this analysis if they had RNA sequencing data, maternal plasma 25(OH)D concentrations measured at enrollment (mid-pregnancy) or delivery, and complete covariate data. All research activities for the CANDLE cohort were approved by the University of Tennessee Health Sciences Center IRB (20) and the ECHO prenatal and early childhood pathways to health consortium (ECHO-PATHWAYS) single IRB (21).

At the mid-pregnancy study enrollment visit, demographic data were collected, including maternal age, race/ethnicity, educational attainment, and health insurance status. Other variables such as pre-pregnancy BMI and Healthy Eating Index (HEI) 2010 were calculated from self-reported data as previously described (22). Neighborhood deprivation index (NDI) was derived with principal components analysis to generate a census tract-level continuous variable incorporating levels of education, professional employment, owner occupied housing, poverty, and unemployment as previously described (23). At the mid-pregnancy visit and late-pregnancy visit (during the third trimester), participant urine samples were collected, which were subsequently used to measure urinary cotinine adjusted for specific gravity as previously described (24). At either urine collection timepoint, urinary cotinine >200 ng/mL was used to classify maternal smoking status as previously described (25,26). At delivery, birth and fetal data were collected, including mode of delivery, labor status, and fetal sex.

### 2.2. Maternal plasma collection and vitamin D measurement

Maternal plasma vitamin D was measured as previously described (27,28). At the mid-pregnancy visit (16-28 weeks’ gestation) and at delivery, maternal blood samples were collected, transported on ice, and centrifuged at 4°C. Aliquots of the resulting plasma were stored at -20°C until further analysis. Samples were processed and frozen within 6 hours of collection. Plasma 25(OH)D concentrations (a total of vitamin D_2_ and D_3_) were measured using a commercial enzymatic immunoassay kit (Immunodiagnostic Systems), according to the manufacturer’s instructions. The analysis was performed at the University of Tennessee Health Science Center in a laboratory that participates in the College of American Pathology Quality Assessment Program for 25(OH)D assays. The minimum detection limit for this assay was 2 ng/mL. National Institute of Standards and Technology SRM972 Vitamin D was used as a standard for quality assurance of 25(OH)D, with interassay variability <6% and precision within 1 SD of mean 25(OH)D concentration.

### 2.3. Placental sample collection and RNA sequencing

The ECHO-PATHWAYS consortium generated placental RNA sequencing data from 794 participants in the CANDLE cohort as described by LeWinn et al (21). Briefly, placental tissue was collected by CANDLE researchers within 15 minutes of delivery and a piece of placental villous tissue approximately 2 cm × 0.5 cm × 0.5 cm was dissected from the middle of the placental parenchyma (26). The tissue was further split into four cubes, which were refrigerated in RNALater at 4°C overnight, transferred to fresh RNALater, and stored at -80°C. The tissue was manually dissected to remove maternal decidual tissue and fetal villous tissue was used for RNA isolation as previously described (26). Briefly, approximately 30 mg of tissue was homogenized using a TissueLyser LT instrument (Qiagen) and RNA was isolated using the AllPrep DNA/RNA/miRNA Universal Kit (Qiagen). Only samples with RNA integrity number >7 as measured using a Bioanalyzer 2100 with RNA 6000 Nanochips (Agilent) were sequenced.

RNA sequencing was performed at University of Washington Northwest Genomics Center as previously described (26). Total RNA was poly-A enriched, complementary DNA libraries were prepared using the TruSeq Stranded mRNA kit (Illumina), and each library was sequenced to an approximate depth of 30 million reads on an Illumina HiSeq 4000 instrument. RNA sequencing quality control was performed using both the FASTX-tool (version 0.0.13) and FastQC (version 0.11.2) toolkits (29). Transcript abundances were estimated by aligning to the GRCh38 transcriptome (Gencode version 33) using Kallisto (30), then collapsed to the gene level using the Bioconductor tximport package (31) and scaled to the average transcript length.

### 2.4. Statistical Analysis of Covariate Data

We used the Institute of Medicine 25(OH)D cutoff for bone health (20 ng/mL) to classify participants as adequate or inadequate (32) and tested the relationship between vitamin D inadequacy and participant characteristics using the Wilcoxon-Mann-Whitney *U* test for continuous variables and Chi-squared tests for categorical variables. We considered *p* < 0.05 significant in these analyses.

### 2.5. Differentially Expressed Gene Identification

RNA sequencing data was filtered to include only protein-coding genes. Expression of these genes was normalized using the trimmed mean of M-values followed by conversion to log counts per million (logCPM) (33). Genes with low expression were removed by filtering for genes with an average logCPM>0 as previously described (26,34,35). Filtering by expression was conducted separately for the samples with plasma 25(OH)D measured at mid-pregnancy and plasma 25(OH)D measured at delivery. After filtering, we tested the association between mid-pregnancy 25(OH)D and expression of 12,892 placental genes and the association between delivery 25(OH)D and expression of 12,893 placental genes. Differentially expressed genes (DEGs) were identified using the *edgeR* limma-voom pipeline (36). Independent linear models were constructed for maternal plasma 25(OH)D at each time point as the exposure variable. We used a directed acyclic graph (DAG) to identify covariates, including confounding and precision variables (**Supplementary Figure 1**). Our independent variable, plasma 25(OH)D concentration, depends on vitamin D intake from foods and supplements, as well as endogenous synthesis.

Vitamin D synthesis is influenced by factors including sun exposure due to latitude, season, clothing, and skin pigmentation (37). While self-identified race may be correlated with skin pigmentation, it is a poor proxy variable (38). Our DAG encodes the assumption that self-reported race, as a social construct, is not related to the placenta transcriptome. Thus, we do not include maternal self-reported race as a confounding variable in this analysis. However, experiencing structural inequities could influence the placenta transcriptome (39). Plasma 25(OH)D concentrations levels have been linked to socioeconomic advantages and higher diet quality in this cohort previously (28,40). Thus, measures of multiple facets related to socioeconomic status and structural inequities, including NDI, education, and health insurance type were identified as confounding variables. These measures are highly correlated with self-reported race in this population. We also identified maternal pre-pregnancy BMI, age, and smoking status as potential confounding variables. Additionally, we identified RNA sequencing batch, delivery method, labor status, and fetal sex as precision variables. The final models were adjusted for NDI, maternal pre-pregnancy BMI, maternal age at delivery, maternal education, maternal health insurance type, smoking status, RNA sequencing batch, delivery method, labor status, and fetal sex.

Our primary analysis of the association between maternal prenatal vitamin D levels and placental gene expression at birth used maternal plasma 25(OH)D at mid-pregnancy as a continuous variable as the exposure of interest. Since vitamin D acts as a hormone, it is possible that responses may be nonlinear (41). Additionally, while a cutoff using bone health as a main endpoint is available (32), there are not universally accepted cutoffs for sufficient vitamin D levels during pregnancy (42,43). Thus, we also used tertiles of plasma 25(OH)D concentrations at mid-pregnancy to evaluate the relationship between vitamin D and placental gene expression as described previously (40). We used the first tertile (from 5.9 to <17.4 ng/mL) as the referent, compared to the second (from 17.4 to <25.1 ng/mL) and third (from 25.1 to 60.2 ng/mL) tertiles of 25(OH)D levels at mid-pregnancy. We also considered that vitamin D levels measured at delivery could influence placental gene expression in samples collected at birth. Thus, we also evaluated the relationship between placental gene expression and plasma 25(OH)D at delivery as both a continuous and a categorical variable. We used the first tertile (from 5.7 to <18.9 ng/mL) as the referent, compared to the second (from 18.9 to <27.0 ng/mL) and third (from 27.0 to 85.0 ng/mL) tertiles of 25(OH)D levels at delivery.

Fetal sex could also modify the relationship between vitamin D levels and placental gene expression, which we tested in sex-stratified models. The interaction between fetal sex and vitamin D levels was assessed after adjustment for the same covariates. We used the Benjamini-Hochberg procedure to control the false discovery rate (FDR) and considered FDR<0.1 significant (44).

### 2.6. Weighted Gene Co-expression Network Analysis

Gene expression may covary, particularly for genes that belong to the same biological pathways or that are regulated by the same transcription factors (TFs), so we conducted Weighted Gene Co-expression Network Analysis (WGCNA) on the entire CANDLE RNA-sequencing data set (*N* = 794). RNA sequencing data was filtered as described above, and count data was then normalized using conditional quantile normalization (*cqn::cqn* function) to gene length and CG composition (45). WGCNA was conducted using the *WGCNA* package (version 1.72-1) (46) as an unsigned network constructed using Pearson’s correlation, hierarchical clustering based on cluster mean averages, and modules containing at least 20 genes. Modules were determined using dynamic tree cut (*WGCNA::cutreeDynamic*).

Modules were characterized by identifying hub genes, conducting KEGG pathway over- representation analysis (*limma::kegga* function) on gene members, and TF over-representation analysis using a placenta-specific transcription regulatory network (47). Genes assigned to a given module that were highly correlated with that module’s eigengene (|*r*|>0.8) were selected as the given module’s hubgenes. In over-representation tests, KEGG pathways and TFs were considered significantly enriched when FDR<0.05. Characterization of all WGCNA modules by hubgenes, over-represented KEGG pathways, and over-represented TFs are presented in **Supplementary Tables 1-3**, respectively.

Multiple linear regression was used to identify WGCNA modules associated with maternal vitamin D levels after adjustment for covariates selected from the DAG. In sex-stratified analyses, the interaction between fetal sex and vitamin D levels was assessed after adjustment for the same covariates. We considered *p*<0.05 significant in these analyses.

## 3. RESULTS

This analysis included *N*=774 CANDLE participants with complete covariate data, RNA sequencing data, and maternal plasma 25(OH)D concentrations at mid-pregnancy and *N*=753 participants with complete covariate data at delivery. Compared to the complete CANDLE cohort (*N*=1503), participants included in this analysis were older, more socioeconomically advantaged, and had higher plasma 25(OH)D concentrations at birth (but not at mid-pregnancy) (**Supplementary Table 4**). The median 25(OH)D concentration at mid-pregnancy was 21.8 ng/mL (IQR: 15.4-26.5 ng/mL) and 324 (41.9%) participants had inadequate levels (**Figure 1A**). At delivery, the median concentration was 23.6 ng/mL (IQR: 16.8-29.1 ng/mL) and 291(38.6%) participants had inadequate levels (**Figure 1B**). Plasma 25(OH)D concentrations between mid-pregnancy and delivery were highly correlated (**Figure 1C**, *N*=752, Spearman’s *ρ*=0.858, *p*<0.001). Participants with adequate plasma 25(OH)D at mid-pregnancy (≥20 ng/mL) were older, had lower pre-pregnancy BMI, lived in neighborhoods with lower NDI scores, had higher HEI-2010 scores, and had higher plasma 25(OH)D concentrations at birth than participants with inadequate plasma 25(OH)D at mid-pregnancy (<20 ng/mL) (**Table 1**). They were also more likely to self-identify as White, have private insurance, have induced labor, and have completed additional education after high school.

**Figure 1.**
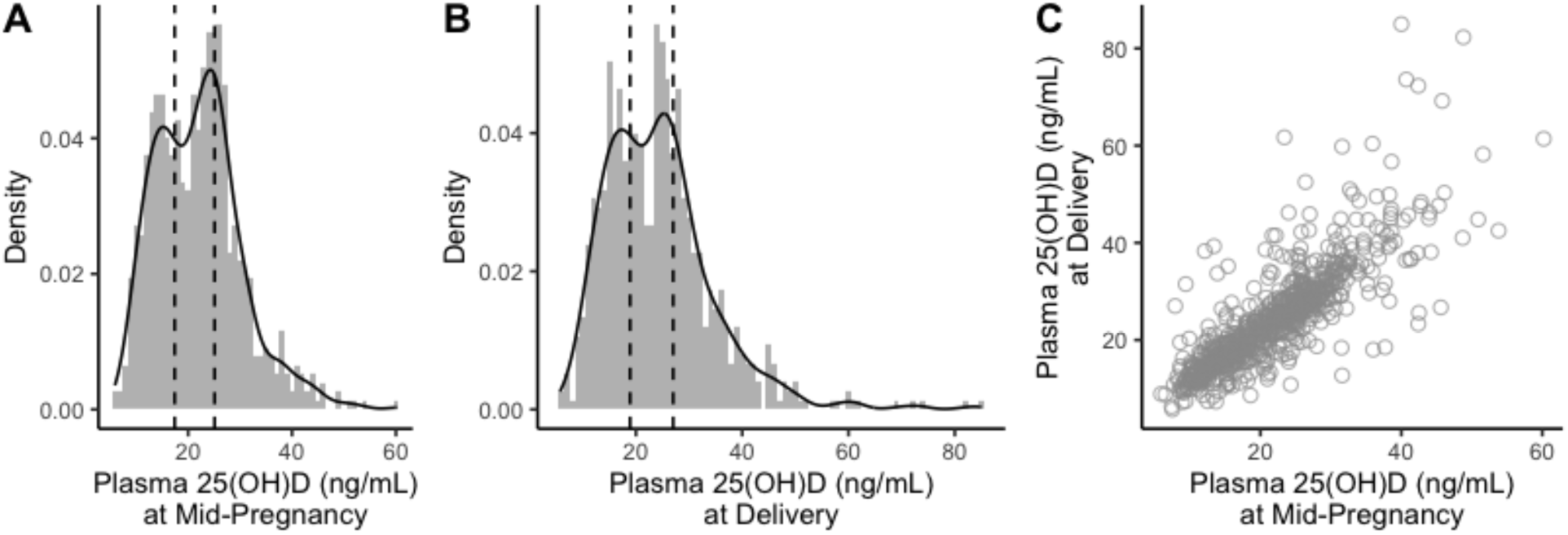
Vitamin D levels at mid-pregnancy and delivery. Density plots of vitamin D levels measured as plasma 25(OH)D levels at (**A**) enrollment during mid-pregnancy (*N*=774) and (**B**) at delivery (*N*=753). Tertile cutoffs are indicated by dotted vertical lines. (**C**) A scatter plot of Vitamin D levels at mid-pregnancy and delivery (*N*=752) shows high correlation.

**Table 1:** Characteristics and socio-demographic factors by maternal vitamin D status at mid-pregnancy.

|  | Inadequate (N=324) | Adequate (N=450) | *p* value |
| --- | --- | --- | --- |
| **Maternal age at birth, years** |  |  | < 0.001 |
| Median | 26 | 28 |  |
| IQR | 22, 30 | 24, 32 |  |
| **Maternal pre-pregnancy BMI** |  |  | 0.002 |
| Median | 27 | 25 |  |
| IQR | 23, 33 | 22, 31 |  |
| **Maternal education** |  |  | < 0.001 |
| < High school | 44 (13.6%) | 20 (4.4%) |  |
| High school graduate/GRE | 168 (51.9%) | 176 (39.1%) |  |
| Graduated college or technical school | 80 (24.7%) | 179 (39.8%) |  |
| Some graduate work or more | 32 (9.9%) | 75 (16.7%) |  |
| **Income, USD** |  |  | < 0.001 |
| Median | 22500 | 50000 |  |
| IQR | 7500, 50000 | 22500, 80000 |  |
| Missing | 31 | 9 |  |
| **Maternal race** |  |  | < 0.001 |
| White | 72 (22.2%) | 222 (49.3%) |  |
| Black/African American | 225 (69.4%) | 208 (46.2%) |  |
| Asian | <10 | <10 |  |
| Multiple race | 24 (7.4%) | 14 (3.1%) |  |
| Other race | <10 | <10 |  |
| **Maternal ethnicity** |  |  | 0.242 |
| Not Hispanic/Latino | 316 (97.5%) | 444 (98.7%) |  |
| Hispanic/Latino | <10 | <10 |  |
| **Fetal sex** |  |  | 0.778 |
| Female | 163 (50.3%) | 231 (51.3%) |  |
| Male | 161 (49.7%) | 219 (48.7%) |  |
| **Labor type** |  |  | 0.048 |
| Spontaneous | 67 (20.7%) | 82 (18.2%) |  |
| Spontaneous, augmented | 104 (32.1%) | 113 (25.1%) |  |
| Induced | 92 (28.4%) | 165 (36.7%) |  |
| No labor | 61 (18.8%) | 90 (20.0%) |  |
| **Delivery method** |  |  | 0.157 |
| Vaginal | 205 (63.3%) | 262 (58.2%) |  |
| C-section | 119 (36.7%) | 188 (41.8%) |  |
| **Cotinine ≥ 200 ng/mL at prenatal visits** |  |  | 0.011 |
| No | 290 (89.5%) | 425 (94.4%) |  |
| Yes | 34 (10.5%) | 25 (5.6%) |  |
| **Gestational age at birth, days** |  |  | 0.123 |
| Median | 274 | 275 |  |
| IQR | 269, 279 | 271, 279 |  |
| Missing | 2 | 2 |  |
| **Neighborhood deprivation index** |  |  | < 0.001 |
| Median | 0.548 | -0.156 |  |
| IQR | -0.344, 1.031 | -0.592, 0.674 |  |
| **Health insurance type** |  |  | < 0.001 |
| Private only | 107 (33.0%) | 276 (61.3%) |  |
| No insurance | <10 | <10 |  |
| Medicaid (TennCare) or Medicare only | 206 (63.6%) | 161 (35.8%) |  |
| TennCare AND Private | 10 (3.1%) | 13 (2.9%) |  |
| **25(OH)D (ng/mL) at mid-pregnancy** |  |  | < 0.001 |
| Median | 14.5 | 25.8 |  |
| IQR | 12.1, 16.9 | 23.1, 29.7 |  |
| **25(OH) D (ng/mL) at delivery** |  |  | < 0.001 |
| Median | 16.2 | 28.1 |  |
| IQR | 13.3, 19.3 | 24.6, 33.7 |  |
| Missing | 7 | 16 |  |
| **HEI-2010, Total Score** |  |  | < 0.001 |
| Median | 59.1 | 63.3 |  |
| IQR | 50.8, 67.4 | 55.3, 71.3 |  |
| Missing | 41 | 32 |  |

Continuous variables are reported as median and interquartile range (IQR), and associations were tested using the Wilcoxon-Mann-Whitney test. Categorical variables are reported as *n* (percentage) and associations were tested using $\chi^2$ test. Maternal vitamin D status at mid-pregnancy was defined as adequate if plasma 25(OH)D ≥ 20 ng/mL or inadequate if plasma 25(OH)D < 20 ng/mL.

We identified 9 DEGs whose placental expression was positively associated and 16 DEGs whose placental expression was inversely associated with plasma 25(OH)D level at mid-pregnancy when controlling for FDR at the 10% level (**Table 2**). The top 5 most upregulated genes were *C1orf21*, *SLC35E2A*, *TRIM44*, *INTS9*, and *DYNC1H1*. The top 5 most down-regulated genes were *COX17*, *AP6V1FNB*, *FAM177A1*, *EMC6*, and *MRPS14*. In sex-stratified analysis (*N* = 394), we identified 1 gene, *COX17*, inversely associated with 25(OH)D among females. Among females, *COX17* expression decreased by 0.74% for each 1 ng/mL increase in mid-pregnancy plasma 25(OH)D concentration (FDR=0.063). We identified no DEGs whose placental expression was associated with 25(OH)D among males (*N*=380).

**Table 2:**
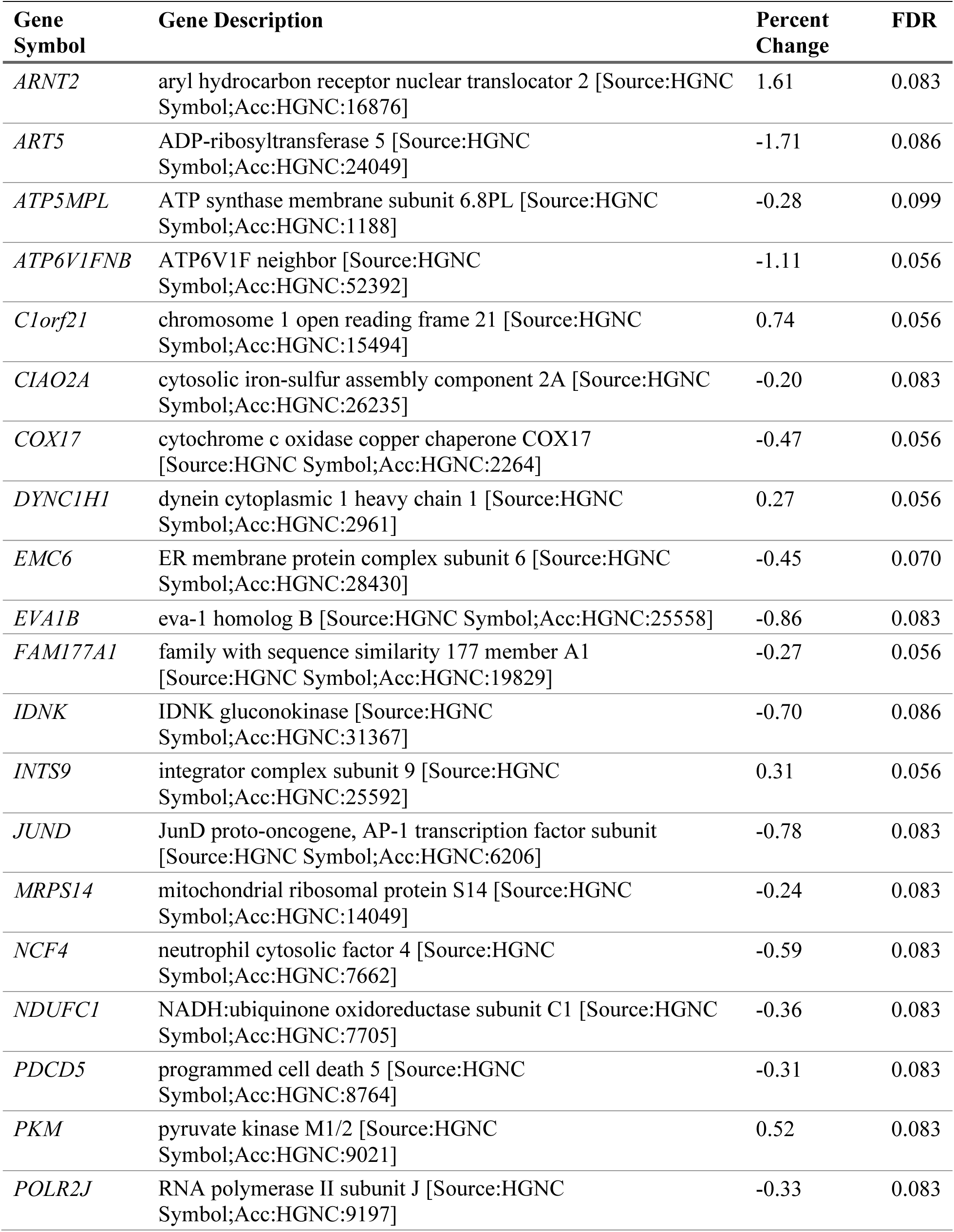
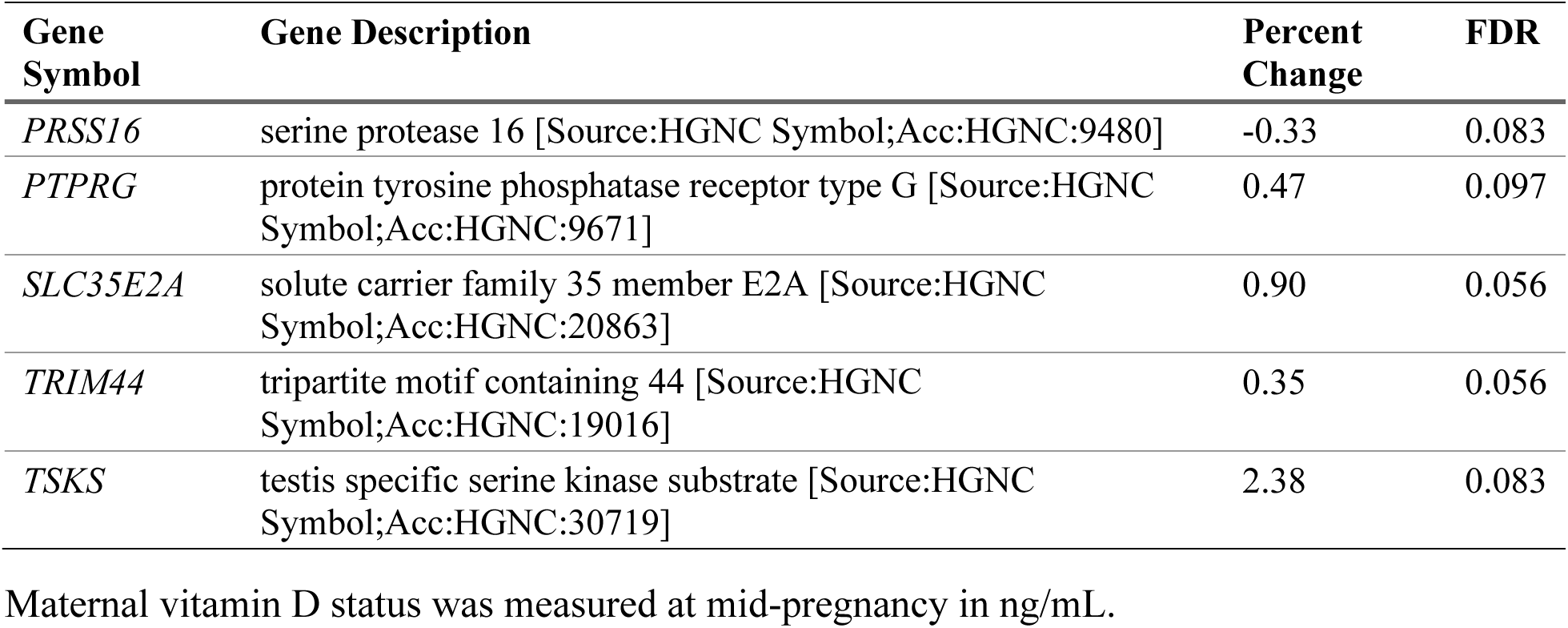
Significant associations between placental gene expression and a 10 ng/mL change in maternal mid-pregnancy vitamin D level (FDR < 0.1)

In the maternal 25(OH)D tertiles and placental gene expression analyses, we identified 44 genes that differed in the second tertile (**Supplementary Table 5**, *n*=261, 17.4–25.0 ng/mL) and 45 genes that differed in the third tertile (**Supplementary Table 6**, *n*=259, 25.1-60.2 ng/mL) compared to the first tertile (*n*=254, 5.9–17.3 ng/mL). The intersection between these DEGs is presented in **Figure 2**. Notably, no DEGs were identified when comparing the second and third tertiles. Two genes that were positively associated with 25(OH)D at mid-pregnancy in our continuous analysis, *ARNT2* and *PKM*, were upregulated in both the second and third tertiles compared to the first (**Figure 2**). Three additional genes that were positively associated with continuous 25(OH)D at mid-pregnancy, *C1orf21*, *DYNC1H1*, *TRIM44*, were upregulated in the third tertile compared to the first (**Figure 2**). There were 7 genes that were inversely associated with continuous 25(OH)D at mid-pregnancy that were also downregulated in the third tertile compared to the first: *COX17*, *EMC6*, *EVA1B*, *FAM177A1*, *JUND*, *MRPS14*, and *NDUFC1* (**Figure 2**).

**Figure 2.**
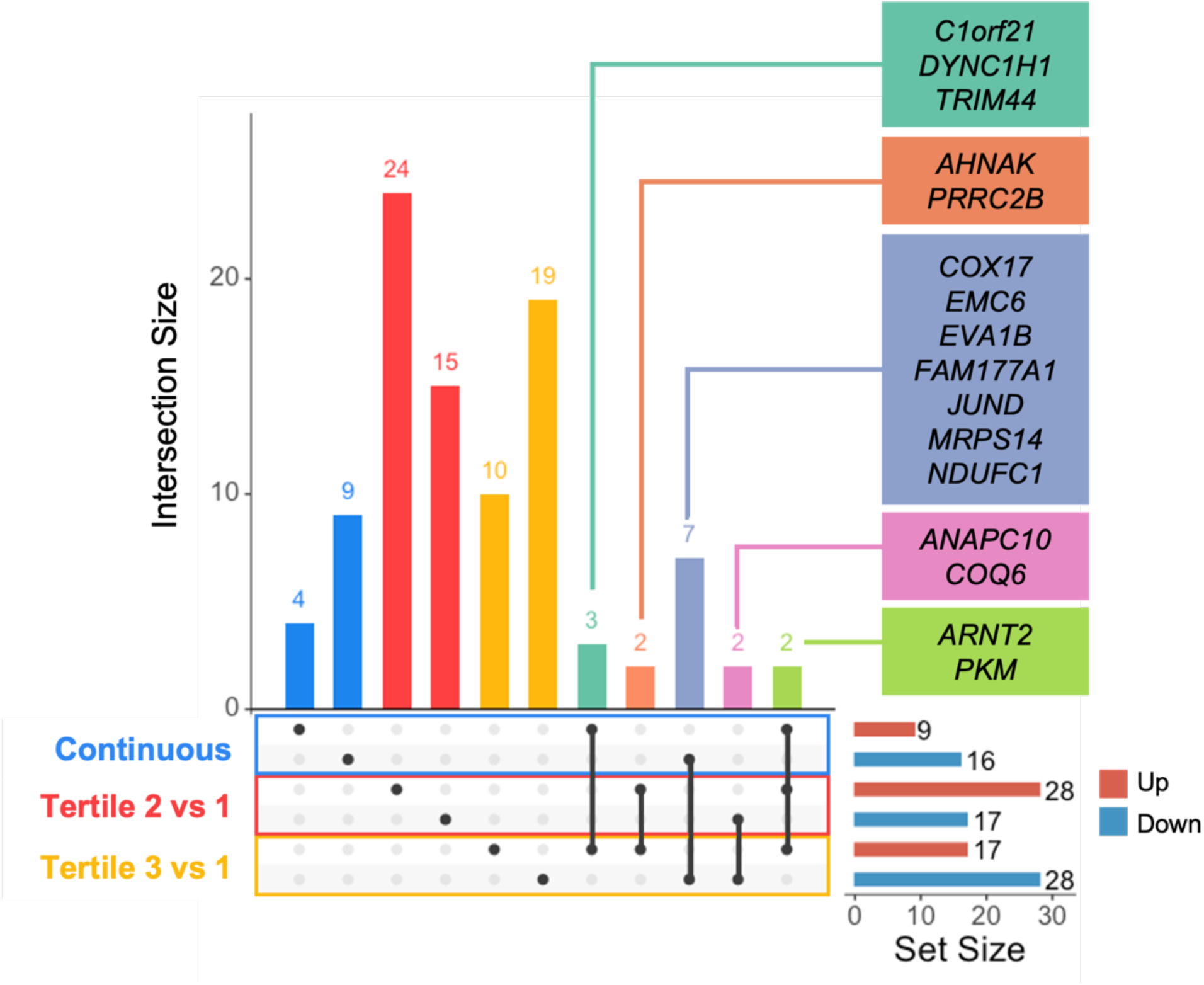
Upset plot depicting shared and distinct changes in placental gene expression. Maternal plasma 25(OH)D levels at mid-pregnancy were analyzed as continuous variable and based on tertile membership. The horizontal bars indicate the total number of upregulated (red) DEGs and downregulated (blue) DEGs in each analysis. The vertical bars indicate the number of unique or overlapping DEGs. The DEGs that are significantly associated with vitamin D levels in more than one analysis are annotated in color-coded boxes.

We also investigated the relationship between maternal 25(OH)D levels, and placental gene expression, both measured at delivery. In our analysis of 25(OH)D as a continuous variable overall and in sex-stratified analysis, we identified no genes whose expression was significantly associated with maternal vitamin D levels at delivery. It was only when we analyzed vitamin D levels as a categorical variable using tertiles that we observed one gene, *MAGEF1*, that was significantly downregulated in the third tertile (logFC=-0.097, percent change from first tertile = -6.52%, FDR=0.012). In female sex-stratified analysis (*N*=384), we identified 1 gene, *CAPN14*, which was most upregulated in the second tertile: *CAPN14* expression was 31.6% higher in the second tertile than the first tertile (logFC=0.396, FDR=0.080) and 24.0% lower in the third tertile when compared to the second tertile (logFC=-0.396, FDR=0.08). We identified no DEGs whose placental expression was associated with 25(OH)D among males (*N*=369).

The relationship between maternal 25(OH)D at mid-pregnancy and delivery and WGCNA modules for placental gene expression were evaluated after adjustment for covariates overall and in sex-stratified analyses (**Figure 3A**). In all samples, the *darkgreen* module was positively associated with maternal 25(OH)D at mid-pregnancy, and the *lightcyan* module was negatively associated with maternal 25(OH)D at both mid-pregnancy and delivery. The *darkgreen* module contained 77 genes, including 15 hubgenes (**Supplementary Table 1**). *MTSS2*, *MICAL3*, *NDRG1*, and *NRIP1* were hubgenes in this module and were also significantly upregulated in the second or third 25(OH)D tertiles compared to the first tertile. The *darkgreen* module genes were not significantly over-represented in any KEGG pathways. However, among the DEGs identified in **Table 2**, the *darkgreen* module was enriched for gene transcription targets of *ARNT2*, *BHLHE40*, *FOSL2*, and *NFKB1,* and *ARNT2* and *BHLHE40* were identified as *darkgreen* module genes (**Figure 3C**, **Supplementary Table 3**). There were 121 genes in the *lightcyan* module, which were significantly over-represented in pathways related to mitochondrial function (oxidative phosphorylation and thermogenesis) and neurotransmission (retrograde endocannabinoid signaling, **Figure 3C**). There were 18 *lightcyan* hub genes, including *EMC6* and *EVA1B*, which were associated with 25(OH)D in our primary model, and *PTRHD1*, which was lower in the third tertile compared to the first tertile. Additionally, *JUND*, another DEG in our primary analysis that was identified as a *lightcyan* module gene, was also identified as TF with gene targets enriched in the *lightcyan* module (**Figure 3B**).

**Figure 3:**
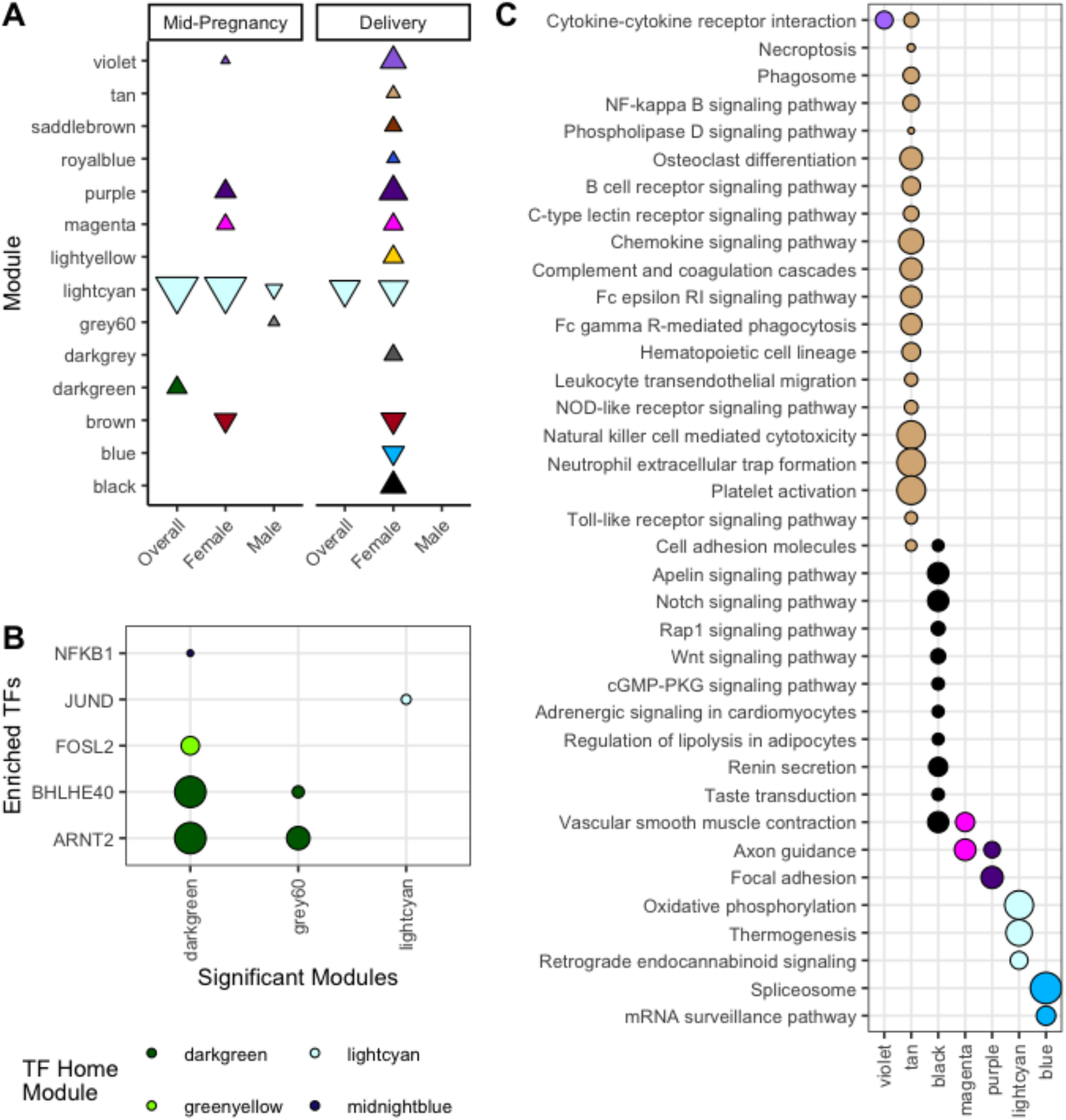
Weighted Gene Co-expression Network Analysis (WGCNA) modules associated with maternal 25(OH)D at mid-pregnancy and delivery. The point size corresponds to -log(*p*) in each panel. (**A**) Associations between modules and maternal 25(OH)D concentrations at mid-pregnancy and delivery were assessed with multiple linear regression adjusted for covariates (*p* < 0.05). Negative associations indicated with downward-pointing triangles and positive associations indicated with upward-pointing triangles. (**B**) Transcription factors (TFs) that were differentially expressed with 25(OH)D and whose gene targets were enriched in these modules were identified using over-representation analysis (FDR < 0.05). (**C**) KEGG pathways that were enriched for module genes were identified through an over-representation analysis (FDR < 0.05).

In males, the *lightcyan* module was negatively associated with maternal 25(OH)D at mid-pregnancy, while the *grey60* (113 genes) were positively associated with maternal 25(OH)D at mid-pregnancy. The *grey60* module did not have DEGs identified as hub genes or enriched KEGG pathways, but the *grey60* module genes were enriched for *ARNT2* gene transcription targets (**Figure 3B**). In females, the *lightcyan* and *brown* (485 genes) modules were negatively associated with maternal 25(OH)D at mid-pregnancy and delivery, while the *blue* (137 genes) module was negatively associated with maternal 25(OH)D at delivery. Also in females, the *magenta* (244 genes), *purple* (232 genes), and *violet* (48 genes) modules were positively associated with maternal 25(OH)D at mid-pregnancy and delivery, while the *black* (127 genes), *darkgrey* (28 genes), *lightyellow* (27 genes), *royalblue* (38 genes), and *tan* (96 genes) modules were positively associated with maternal 25(OH)D at delivery (**Figure 3A**). The *black* module was enriched for pathways related to vascular function, including angiogenesis (Apelin, Notch, and Wnt signaling) and vascular smooth muscle contraction. The *blue* module was enriched for mRNA surveillance and the spliceosome pathway. The *magenta* module was enriched for vascular smooth muscle contraction and axon guidance. The *purple* module was enriched for focal adhesion and axon guidance. The *violet* module was enriched for cytokine-cytokine receptor interaction. The tan module was enriched for 20 pathways related to immune responses, including chemokine signaling, natural killer cell mediated cytotoxicity, and platelet activation (**Figure 3C**). The *black*, *blue*, *magenta*, *purple*, and *violet* modules did not have DEGs identified in **Table 2** or **Supplementary Tables 5-6** among their hubgenes or enriched TFs.

## 4. DISCUSSION

This analysis is the first to investigate the relationship between prenatal vitamin D levels quantified at different time points in pregnancy and the placental transcriptome in a prospective pregnancy cohort. In our analysis, we identified 25 placental genes whose expression at birth was associated with maternal vitamin D levels during mid-pregnancy but not with maternal vitamin D levels at delivery. We identified an additional 72 genes whose expression was altered in the second and/or third tertiles of maternal mid-pregnancy vitamin D levels compared to the lowest tertile, including *AHNAK*, *ANAPC10*, *ARNT2*, *COQ6*, *PKM*, and *PRRC2B*, which were significantly different from the lowest quartile in both the second and third tertiles. All together, these DEGs encode proteins related to energy metabolism, cytoskeletal dynamics, and placental transcriptional regulation. This suggests that maternal 25(OH)D levels during pregnancy could play a role in programming placental biology. We also conducted WGCNA to investigate associations between gene modules that are co-expressed and co-regulated and vitamin D levels. We identified 7 modules related to maternal vitamin D levels at mid-pregnancy and 12 modules related to maternal vitamin D levels at delivery, which provided further support for associations between vitamin D levels and mitochondrial function, cytoskeletal function, and key TFs.

The placenta undergoes rapid development from implantation through early pregnancy and continues to grow and build increasingly complex vasculature throughout gestation (48,49). As the placenta grows, its energetic needs increase (50). From an evolutionary perspective, one of the first roles of 1,25(OH)_2_D mediated by the VDR was regulating energy metabolism (51).

Indeed, in this study, vitamin D levels were positively associated with expression of genes encoding glycolytic enzymes (*ALDOA*, *PKM*), but inversely associated with mitochondrial proteins (*MPC2*, *MRPS14*, *TIMM17A*) and specifically components of the electron transport chain (*ATP5MPL*, *COX6C*, *COX8A*, *COX17*, *NDUFB6*, *NDUFC1*). *PKM* encodes pyruvate kinase, a regulator of trophoblast invasion in the placenta. 1,25(OH)_2_D has been shown to increase pyruvate kinase activity in human fibroblast and neuroblastoma cell lines (52–54). *In vitro* studies suggest that 1,25(OH)_2_D and VDR regulate mitochondrial function in skeletal muscle (55–57) and vitamin D supplementation improved strength and muscle mass in older adults (57). In human skeletal muscle cells, 1,25(OH)_2_D treatment resulted in extensive changes in expression of genes related not only to mitochondrial function (including downregulated *COX17* expression, as we report here), but also cytoskeletal and intracellular membrane trafficking (55).

We also report associations between maternal vitamin D levels and placental expression of genes involved in cytoskeletal function: motor proteins (*DYNC1H1*, *MYH9*), microtubule-associated proteins (*CLIP2*, *DCTN2*, *GABARAPL2*, *MAP1LC3A*, *MICAL3*), and other genes related to cellular transport (*AHNAK*, *DSP*, *FLNB*, *JPH2*, *MINK1*, *MTSS2*, *RABIF*, *RHOV*, *TANC2*). Some of these genes are essential for placental development, such as *MYH9*, which is required for placental vascular development and knockouts lead to embryonic death (58,59). Additionally, the protein encoded by *MAP1LC3A* also binds to the VDR and promotes its translocation to the nucleus (60). The cytoskeleton is crucial in key placental processes, including trophoblast invasion, cellular division and proliferation, autophagy, and intracellular transport (61). Vitamin D regulates cytoskeletal reorganization in other tissues (62,63). In trophoblasts, vitamin D treatment promotes cellular migration and invasion in vitro, processes that are dependent on cytoskeletal reorganization (11,12,64,65). Cytoskeletal proteins and intracellular membrane trafficking pathways are also necessary for placental nutrient transport, including trafficking amino acid and glucose transporters, as well as fusion into the multinucleate syncytium (66,67). Thus, cytoskeletal and trafficking related DEGs support the importance of prenatal vitamin D across domains of placental function.

As a ligand for the VDR, 1,25(OH)_2_D is involved in regulating transcription of VDR response genes and we observe associations between vitamin D levels and genes related to the machinery of transcription (*POLR2J*), RNA processing (*GCFC2*, *INTS9*, *SF1*) and other TFs (*ARNT2*, *BHLHE40*, *FOSL2*, *JUND*, *NFKB1*) (68,69). Indeed, the VDR is also thought to influence the transcription of other TFs, leading to a two-phase response to vitamin D (70). Thus, we also investigated gene coexpression using WGCNA to identify modules of coexpressed genes associated with 25(OH)D levels. We identified 2 modules associated with 25(OH)D levels: the *lightcyan* modules that was inversely associated with 25(OH)D levels at mid-pregnancy and delivery and the *darkgreen* modules that was positively associated with 25(OH)D levels at mid-pregnancy. In this analysis, *ARNT2*, *BHLHE40*, *FOSL2*, and *NFKB1* were positively associated with vitamin D levels and their targets were enriched in the *darkgreen* WGCNA module. In a placental-specific transcription regulatory network, *BHLHE40*, *FOSL2*, and *NFKB1* regulate each other (47). Additionally, *ARNT2* is regulated by *VDR* and regulates *VDR* and *BHLHE40* (47). *ARNT2* and its paralog *ARNT* dimerize with *HIF1A* to regulate the hypoxic response and dimerize with *AHR* to regulate placental vascular development through pregnancy (68,71). *NFKB1* encodes multiple gene products that mediate both inflammatory and anti-inflammatory gene expression (72). Vitamin D treatment in naïve B cells resulted in reduced *NFKB1* mRNA expression (73), though we observed elevated *NFKB1* expression in placental samples here. While elevated expression of placental *NFKB1* has been linked to preeclampsia and preterm labor, expression is also elevated in term labored placentas compared to unlabored controls, suggesting a physiological role in parturition (74–76). More work is needed to elucidate these molecular mechanisms.

*FOSL2* and *JUND* are two subunits of the activator protein-1 (AP-1) transcription factor, which plays an essential role in development as a regulator of cellular differentiation, proliferation, and apoptosis (77). AP-1 is made up of protein subunits from the JUN family and the FOS family and the specific composition of the AP-1 dimer influences its function (77). *FOSL2* is crucial in bone formation and activated by 1,25(OH)_2_D in osteoclasts (78). While other members of the FOS family are necessary for trophoblast invasion, migration, and development and *FOSL2* is highly expressed in extravillous trophoblasts, the role of *FOSL2* in the placenta is less clear. We report that *JUND* is negatively associated with vitamin D and its targets are enriched in the *lightcyan* module. *JUND* is also activated by 1,25(OH)_2_D in some cells, including osteoclasts (multinucleated cells involved in dissolving the extracellular matrix in bone resorption), but its expression was not affected in bone-forming osteoblasts (78,79). In the placenta, *JUND* is linked to trophoblast syncytialization (80,81).

In this study, we also investigated fetal sex as an effect modifier for the relationship between maternal 25(OH)D levels and placental gene expression. We found evidence for effect modification by sex only for the association between mid-pregnancy 25(OH)D levels and COX17 and delivery 25(OH)D tertile membership and *CAPN14*. Thus, at the level of individual genes, we did not find evidence of broad effects of sex on the association between vitamin D levels and placental gene expression. However, when investigating associations between vitamin D levels and patterns of gene coexpression, identified 4 modules that were associated with vitamin D levels at mid-pregnancy only in females and one module that was associated with vitamin D levels at mid-pregnancy only in males. The *magenta* and *purple* modules were related to aspects of cytoskeletal function and the *violet* module was related to cellular signaling, in keeping with roles of vitamin D we have previously discussed. At delivery, the *lightcyan* module was significantly negatively associated with vitamin D levels overall and in females, but not males. Additionally, vitamin D levels at delivery were associated with 11 other modules only in females, including the *tan* (immune function) and *black* (vascular function). These patterns only observed in females might be related to sex differences in vitamin D metabolism (82). Evidence from mice shows male placentas express higher levels of vitamin D-inactivating *Cyp24a1* in vitamin D sufficiency and lower levels of vitamin D-activating *Cyp27b1* in vitamin D deficiency compared to equivalent female placentas (18). In human trophoblasts, testosterone treatment downregulated *CYP27B1* expression and upregulated *CYP24A1* expression in a time- and dose-dependent manner and in term placentas, males had lower *CYP27B1* mRNA expression (19). Further study is needed to investigate associations between sex differences in vitamin D homeostasis and placental gene expression.

Our findings align with the handful of studies that have investigated the effect of vitamin D on the placental transcriptome using RNA sequencing (10,83). In an *in vitro* study using cultured primary trophoblasts treated with 20 μM 25(OH)D for 8 hours and reported upregulation of *MGAT3*, *MICAL*, and *TANC2* (10) in agreement with our findings reported here. Additionally, downregulation of immune, inflammatory, and cytokine-binding pathways in conjunction with upregulation of transcriptional regulatory pathways were reported (10). Combining transcriptomic and proteomic analysis also identified cytoskeletal binding as an enriched pathway following 25(OH)D treatment (10). Recently, a randomized controlled trial providing the recommended dose of 10 μg (400 international units [IU]) vitamin D/day or the high dose of 90 μg (3,600 IU) vitamin D/day from the late first trimester through delivery employed RNA sequencing to investigate placental gene expression in a random subset of the study (*N* = 70) (83). This study reported that high-dose vitamin D supplementation was associated with enrichment in the cell adhesion pathway and downregulated expression with *JPH1* (83). While *JPH1* was not in our filtered dataset, we observed that its paralog *JPH2* was downregulated in the third tertile compared to the first tertile of vitamin D status at mid-pregnancy. Additionally, we report that known targets of the *VDR* or 1,25(OH)_2_D treatment in other cell types were upregulated with higher prenatal 25(OH)D levels, including *BHLHE40*, *DIO2*, *NRIP1*, *PIK3R1*, and *TIMP3* (84–88). DEGs associated with 25(OH)D as a continuous or categorical variable in this analysis should be experimentally validated using a range of doses in future studies.

Our results should be interpreted in the context of limitations. First, we conducted RNA sequencing on placental samples collected at birth, which provides a snapshot of a highly coordinated temporal process. Thus, variation in gene expression at birth may not reflect placental gene expression during gestation. Second, expression was quantified in bulk samples and could mask cell type-specific changes in gene expression related to vitamin D levels (89). The placenta is a complex organ made up of multiple cell types, including trophoblasts, fibroblasts, endothelial cells, and immune cells, which may respond to vitamin D in a cell type-specific manner. Additionally, given the observational study design, we cannot draw causal links between mid-pregnancy vitamin D levels and placental gene expression and there is still the possibility of residual confounding, despite adjustment for a variety of covariates. This study has several strengths. First, because all participants were recruited from the same county, potential confounding by geographical differences in sunlight exposure was reduced. Second, this richly characterized and socioeconomically and racially diverse cohort allowed us to conduct the largest analysis to date of prenatal vitamin D levels and placental gene expression. Third, this analysis employed a transcriptome-wide approach, which enabled us to holistically assess how vitamin D might influence placental function.

These findings in a large, diverse, prospective cohort study identify roles for vitamin D in placental energy metabolism, cytoskeletal function, and transcriptional regulation. Notably, prenatal vitamin D levels during mid-pregnancy were related to placental expression of 97 DEGs, while vitamin D levels measured at delivery were associated with only 2 DEGs. This suggests that mid-pregnancy vitamin D could play a role in programing placental gene expression. Among these DEGs, we identified important transcriptional regulators. Our findings also suggest that sex-specific effects of vitamin D may be subtle and were mainly observed in the co-expression analysis. Future research is needed to investigate the potential programing effect of vitamin D during pregnancy on placental mitochondrial metabolism, intracellular transport, and transcriptional regulation.

## ACKNOWLEDGEMENTS

We would like to thank the families for their participation in the CANDLE study. We would also like to thank the study staff, data teams, and investigators involved in the CANDLE cohort and ECHO PATHWAYS consortium for their contributions. This manuscript has been reviewed by PATHWAYS for scientific content and consistency of data interpretation with previous PATHWAYS publications.

## ABBREVIATIONS

1,25(OH)2D: 1,25-dihydroxyvitamin D
25(OH)D: 25-hydroxyvitamin D
CANDLE: Conditions Affecting Neurocognitive Development and Learning in Early childhood
DEG: Differentially expressed gene
FDR: False discovery rate
HEI-2010: Healthy Eating Index 2010
IU: International units
NDI: Neighborhood Deprivation Index
TF: Transcription factor
WGCNA: Weighted gene co-expression network analysis

