## Supplement 2 for "Associations Between Prenatal Vitamin D and Placental Gene Expression"

[1] Center for Developmental Biology and Regenerative Medicine, Seattle Children's Research Institute, Seattle, WA, United States
[2] Department of Health Behavior and Nutrition Sciences, University of Delaware, Newark, DE, United States
[3] Department of Environmental and Occupational Health Sciences, University of Washington, Seattle, WA, United States
[4] Department of Preventive Medicine, University of Tennessee Health Sciences Center, Memphis, TN, United States
[5] Department of Pharmacology, Addiction Science, and Toxicology, University of Tennessee Health Sciences Center, Memphis, TN, United States
[6] Department of Epidemiology, University of Washington, Seattle, WA, United States
[7] Icahn School of Medicine at Mount Sinai, New York, NY, United States
[8] Department of Psychiatry and Behavioral Sciences, University of California, San Francisco, San Francisco, CA, United States
[9] Department of Pediatrics, University of California, San Francisco, San Francisco, CA, United States
[10] Center for Child Health, Behavior, and Development, Seattle Children's Research Institute, Seattle, WA, United States
[11] Department of Pediatrics, University of Washington, Seattle, WA, United States

### CONTENTS

**Supplementary Figure 1**. Directed Acyclic Graph

**Supplementary Table 4**: Characteristics and socio-demographic factors for CANDLE participants included in this analysis compared to other CANDLE participants

**Supplementary Table 5**: Significant associations between the second tertile of maternal prenatal vitamin D concentrations and placental gene expression (FDR < 0.1) using the first tertile as a reference

**Supplementary Table 6**: Significant associations between the third tertile of maternal prenatal 25(OH)D concentrations and placental gene expression (FDR < 0.1) using the first tertile as a reference

* **Supplementary Tables 1-3** can been found in Supplement 2.

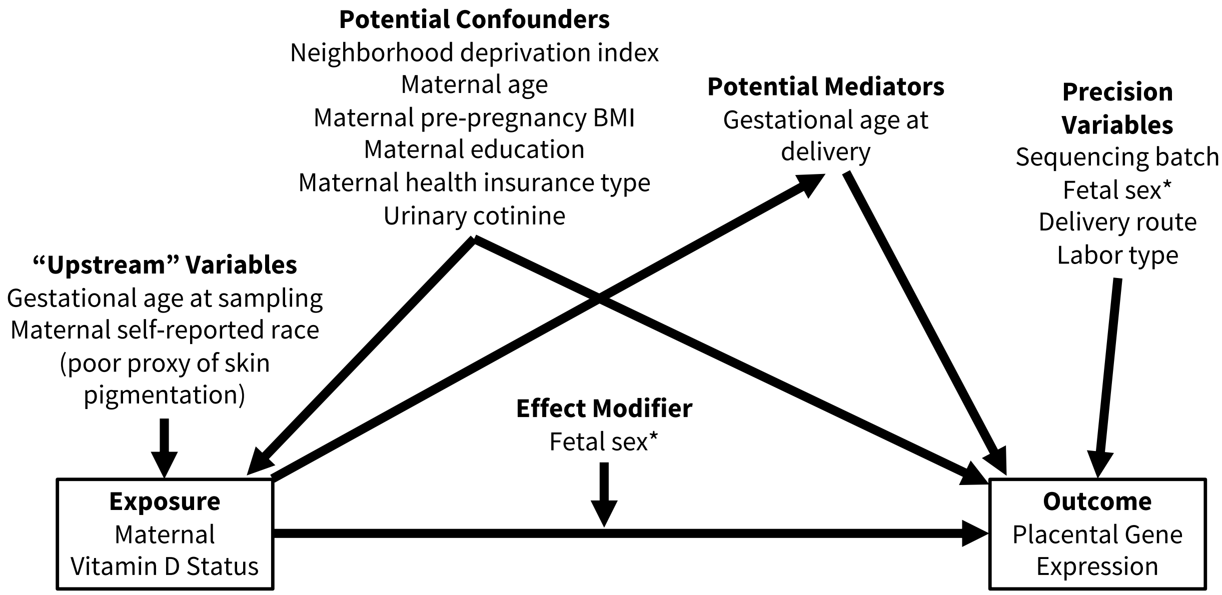


**Supplemental Figure 1**. Directed Acyclic Graph (DAG). * Fetal sex is treated as a precision variable in the overall model and as an effect modifier in the sex-stratified models.

**Supplementary Table 4**: Characteristics and socio-demographic factors for CANDLE participants included in this analysis compared to other CANDLE participants.

| | Excluded (*N*=729) | Included (*N*=774) | *p* value |
|---|---|---|---|
| **Maternal age at birth, years** | | | < 0.001 |
| Median | 25 | 27 | |
| IQR | 21, 29 | 23, 31 | |
| *Missing* | 25 | 0 | |
| **Maternal pre-pregnancy BMI** | | | 0.045 |
| Median | 25 | 26 | |
| IQR | 22, 31 | 22, 32 | |
| *Missing* | 5 | 0 | |
| **Maternal education** | | | < 0.001 |
| < High School | 120 (16.5%) | 64 (8.3%) | |
| High school graduate/GRE | 365 (50.2%) | 344 (44.4%) | |
| Graduated college or technical school | 178 (24.5%) | 259 (33.5%) | |
| Some graduate work or more | 64 (8.8%) | 107 (13.8%) | |
| *Missing* | 2 | 0 | |
| **Income** | | | < 0.001 |
| Median | 22500 | 40000 | |
| IQR | 7500, 50000 | 17500, 70000 | |
| *Missing* | 95 | 40 | |
| **Maternal race** | | | < 0.001 |
| White | 173 (23.8%) | 294 (38.0%) | |
| Black/African American | 503 (69.2%) | 433 (55.9%) | |
| Asian | <10 | <10 | |
| Native Hawaiian/Other Pacific Islander | <10 | <10 | |
| American Indian/Alaska Native | <10 | <10 | |
| Multiple race | 39 (5.4%) | 38 (4.9%) | |
| Other race | <10 | <10 | |
| *Missing* | 2 | 0 | |
| **Maternal ethnicity** | | | 0.288 |
| Not Hispanic/Latino | 708 (97.4%) | 760 (98.2%) | |
| Hispanic/Latino | 19 (2.6%) | 14 (1.8%) | |
| *Missing* | 2 | 0 | |
| **Fetal sex** | | | 0.326 |
| Female | 333 (48.3%) | 394 (50.9%) | |
| Male | 356 (51.7%) | 380 (49.1%) | |
| *Missing* | 40 | 0 | |
| **Labor type** | | | < 0.001 |
| Spontaneous | 191 (28.0%) | 149 (19.3%) | |
| Spontaneous, augmented | 201 (29.5%) | 217 (28.0%) | |

| | Excluded (*N*=729) | Included (*N*=774) | *p* value |
|---|---|---|---|
| Induced | 191 (28.0%) | 257 (33.2%) | |
| No labor | 98 (14.4%) | 151 (19.5%) | |
| *Missing* | 48 | 0 | |
| **Delivery method** | | | 0.030 |
| Vaginal | 449 (65.8%) | 467 (60.3%) | |
| C-section | 233 (34.2%) | 307 (39.7%) | |
| *Missing* | 47 | 0 | |
| **Cotinine ≥ 200 ng/mL at prenatal visits** | | | < 0.001 |
| No | 553 (86.0%) | 715 (92.4%) | |
| Yes | 90 (14.0%) | 59 (7.6%) | |
| *Missing* | 86 | 0 | |
| **Gestational age at birth, days** | | | 0.119 |
| Median | 274 | 275 | |
| IQR | 266, 280 | 270, 279 | |
| *Missing* | 73 | 4 | |
| **NDI** | | | < 0.001 |
| Median | 0.531 | 0.066 | |
| IQR | -0.305, 1.186 | -0.489, 0.870 | |
| *Missing* | 76 | 0 | |
| **Health insurance type** | | | < 0.001 |
| No insurance | <10 | <10 | |
| Medicaid (TennCare) or Medicare only | 492 (67.5%) | 367 (47.4%) | |
| TennCare AND Private | 19 (2.6%) | 23 (3.0%) | |
| Private only | 217 (29.8%) | 383 (49.5%) | |
| **Vitamin D levels (ng/mL) at mid-pregnancy** | | | 0.048 |
| Median | 19.8 | 21.8 | |
| IQR | 14.5, 26.9 | 15.4, 26.5 | |
| *Missing* | 26 | 0 | |
| **Vitamin D status (ng/mL) at birth** | | | < 0.001 |
| Median | 19.1 | 23.6 | |
| IQR | 14.4, 26.4 | 16.8, 29.1 | |
| *Missing* | 302 | 23 | |
| **HEI-2010, Total Score** | | | < 0.001 |
| Median | 58.7 | 62.0 | |
| IQR | 51.0, 67.0 | 52.8, 69.4 | |
| *Missing* | 108 | 73 | |

Continuous variables are reported as median and interquartile range (IQR) and associations were tested using the Wilcoxon-Mann-Whitney test. Categorical variables are reported as *n* (percentage) and associations were tested using $\chi^2$ test.

**Supplementary Table 5**: Significant associations between the second tertile of maternal prenatal vitamin D concentrations and placental gene expression (FDR < 0.1) using the first tertile as a reference

| Gene Symbol | Gene Description | Percent Change | FDR |
|---|---|---|---|
| *AHNAK* | AHNAK nucleoprotein [Source:HGNC Symbol;Acc:HGNC:347] | 13.34 | 0.013 |
| *ALDOA* | aldolase, fructose-bisphosphate A [Source:HGNC Symbol;Acc:HGNC:414] | 8.76 | 0.073 |
| *ANAPC10* | anaphase promoting complex subunit 10 [Source:HGNC Symbol;Acc:HGNC:24077] | -5.71 | 0.071 |
| *ARNT2* * | aryl hydrocarbon receptor nuclear translocator 2 [Source:HGNC Symbol;Acc:HGNC:16876] | 55.95 | 0.012 |
| *BHLHE40* | basic helix-loop-helix family member e40 [Source:HGNC Symbol;Acc:HGNC:1046] | 25.85 | 0.098 |
| *BTG2* | BTG anti-proliferation factor 2 [Source:HGNC Symbol;Acc:HGNC:1131] | -15.88 | 0.012 |
| *C18orf21* | chromosome 18 open reading frame 21 [Source:HGNC Symbol;Acc:HGNC:28802] | -5.01 | 0.073 |
| *CHST11* | carbohydrate sulfotransferase 11 [Source:HGNC Symbol;Acc:HGNC:17422] | -7.59 | 0.073 |
| *CITED2* | Cbp/p300 interacting transactivator with Glu/Asp rich carboxy-terminal domain 2 [Source:HGNC Symbol;Acc:HGNC:1987] | 22.43 | 0.098 |
| *CLIP2* | CAP-Gly domain containing linker protein 2 [Source:HGNC Symbol;Acc:HGNC:2586] | 14.98 | 0.073 |
| *COQ6* | coenzyme Q6, monooxygenase [Source:HGNC Symbol;Acc:HGNC:20233] | -6.97 | 0.073 |
| *DIO2* | iodothyronine deiodinase 2 [Source:HGNC Symbol;Acc:HGNC:2884] | 71.37 | 0.098 |
| *DSP* | desmoplakin [Source:HGNC Symbol;Acc:HGNC:3052] | 11.11 | 0.088 |
| *ERCC3* | ERCC excision repair 3, TFIIH core complex helicase subunit [Source:HGNC Symbol;Acc:HGNC:3435] | -4.07 | 0.098 |
| *ERGIC1* | endoplasmic reticulum-golgi intermediate compartment 1 [Source:HGNC Symbol;Acc:HGNC:29205] | 10.19 | 0.099 |
| *FLNB* | filamin B [Source:HGNC Symbol;Acc:HGNC:3755] | 20.65 | 0.098 |
| *FOSL2* | FOS like 2, AP-1 transcription factor subunit [Source:HGNC Symbol;Acc:HGNC:3798] | 20.56 | 0.019 |
| *GGH* | gamma-glutamyl hydrolase [Source:HGNC Symbol;Acc:HGNC:4248] | -11.01 | 0.086 |
| *GOT1* | glutamic-oxaloacetic transaminase 1 [Source:HGNC Symbol;Acc:HGNC:4432] | -9.63 | 0.098 |
| *HUWE1* | HECT, UBA and WWE domain containing E3 ubiquitin protein ligase 1 [Source:HGNC Symbol;Acc:HGNC:30892] | 5.13 | 0.053 |
| *KMO* | kynurenine 3-monooxygenase [Source:HGNC Symbol;Acc:HGNC:6381] | -17.29 | 0.071 |
| *MGAT3* | beta-1,4-mannosyl-glycoprotein 4-beta-N-acetylglucosaminyltransferase [Source:HGNC Symbol;Acc:HGNC:7046] | 15.34 | 0.099 |
| *MMADHC* | metabolism of cobalamin associated D [Source:HGNC Symbol;Acc:HGNC:25221] | -3.62 | 0.098 |
| *MMUT* | methylmalonyl-CoA mutase [Source:HGNC Symbol;Acc:HGNC:7526] | -5.84 | 0.071 |

| Gene Symbol | Gene Description | Percent Change | FDR |
|---|---|---|---|
| *MTRES1* | mitochondrial transcription rescue factor 1 [Source:HGNC Symbol;Acc:HGNC:17971] | -5.31 | 0.073 |
| *MTSS2* | MTSS I-BAR domain containing 2 [Source:HGNC Symbol;Acc:HGNC:25094] | 16.30 | 0.073 |
| *NDRG1* | N-myc downstream regulated 1 [Source:HGNC Symbol;Acc:HGNC:7679] | 20.14 | 0.098 |
| *NPR3* | natriuretic peptide receptor 3 [Source:HGNC Symbol;Acc:HGNC:7945] | 24.66 | 0.073 |
| *NRIP1* | nuclear receptor interacting protein 1 [Source:HGNC Symbol;Acc:HGNC:8001] | 21.46 | 0.095 |
| *PIK3R1* | phosphoinositide-3-kinase regulatory subunit 1 [Source:HGNC Symbol;Acc:HGNC:8979] | 12.99 | 0.095 |
| *PKM* * | pyruvate kinase M1/2 [Source:HGNC Symbol;Acc:HGNC:9021] | 12.11 | 0.024 |
| *PPP1R12C* | protein phosphatase 1 regulatory subunit 12C [Source:HGNC Symbol;Acc:HGNC:14947] | 9.85 | 0.098 |
| *PRRC2B* | proline rich coiled-coil 2B [Source:HGNC Symbol;Acc:HGNC:28121] | 7.84 | 0.098 |
| *RABIF* | RAB interacting factor [Source:HGNC Symbol;Acc:HGNC:9797] | -4.82 | 0.031 |
| *RHOV* | ras homolog family member V [Source:HGNC Symbol;Acc:HGNC:18313] | -15.19 | 0.068 |
| *SLC4A1AP* | solute carrier family 4 member 1 adaptor protein [Source:HGNC Symbol;Acc:HGNC:13813] | -4.86 | 0.053 |
| *SLC6A8* | solute carrier family 6 member 8 [Source:HGNC Symbol;Acc:HGNC:11055] | 12.89 | 0.073 |
| *SUPT5H* | SPT5 homolog, DSIF elongation factor subunit [Source:HGNC Symbol;Acc:HGNC:11469] | 4.90 | 0.073 |
| *TAF8* | TATA-box binding protein associated factor 8 [Source:HGNC Symbol;Acc:HGNC:17300] | -5.10 | 0.098 |
| *TANC2* | tetratricopeptide repeat, ankyrin repeat and coiled-coil containing 2 [Source:HGNC Symbol;Acc:HGNC:30212] | 16.99 | 0.026 |
| *TIMP3* | TIMP metallopeptidase inhibitor 3 [Source:HGNC Symbol;Acc:HGNC:11822] | 25.92 | 0.098 |
| *UPK1B* | uroplakin 1B [Source:HGNC Symbol;Acc:HGNC:12578] | 42.44 | 0.078 |
| *ZFP36L1* | ZFP36 ring finger protein like 1 [Source:HGNC Symbol;Acc:HGNC:1107] | 10.58 | 0.053 |
| *ZNF395* | zinc finger protein 395 [Source:HGNC Symbol;Acc:HGNC:18737] | 10.15 | 0.086 |

* indicates a gene also identified as a DEG in the primary analysis.

**Supplementary Table 6**: Significant associations between the third tertile of maternal prenatal 25(OH)D concentrations and placental gene expression (FDR < 0.1) using the first tertile as a reference

| Gene Symbol | Gene Description | Percent Change | FDR |
|---|---|---|---|
| *AHNAK* | AHNAK nucleoprotein [Source:HGNC Symbol;Acc:HGNC:347] | 11.37 | 0.078 |
| *ANAPC10* | anaphase promoting complex subunit 10 [Source:HGNC Symbol;Acc:HGNC:24077] | -6.18 | 0.042 |
| *ARNT2* * | aryl hydrocarbon receptor nuclear translocator 2 [Source:HGNC Symbol;Acc:HGNC:16876] | 53.06 | 0.019 |
| *C1orf21* * | chromosome 1 open reading frame 21 [Source:HGNC Symbol;Acc:HGNC:15494] | 14.69 | 0.062 |
| *CEBPZOS* | CEBPZ opposite strand [Source:HGNC Symbol;Acc:HGNC:49288] | -5.32 | 0.062 |
| *COQ6* | coenzyme Q6, monooxygenase [Source:HGNC Symbol;Acc:HGNC:20233] | -6.99 | 0.079 |
| *COX17* * | cytochrome c oxidase copper chaperone COX17 [Source:HGNC Symbol;Acc:HGNC:2264] | -9.32 | 0.033 |
| *COX6C* | cytochrome c oxidase subunit 6C [Source:HGNC Symbol;Acc:HGNC:2285] | -6.82 | 0.100 |
| *COX8A* | cytochrome c oxidase subunit 8A [Source:HGNC Symbol;Acc:HGNC:2294] | -7.35 | 0.078 |
| *DCTN2* | dynactin subunit 2 [Source:HGNC Symbol;Acc:HGNC:2712] | -4.08 | 0.094 |
| *DIP2C* | disco interacting protein 2 homolog C [Source:HGNC Symbol;Acc:HGNC:29150] | 15.30 | 0.089 |
| *DYNC1H1* * | dynein cytoplasmic 1 heavy chain 1 [Source:HGNC Symbol;Acc:HGNC:2961] | 6.13 | 0.019 |
| *EMC6* * | ER membrane protein complex subunit 6 [Source:HGNC Symbol;Acc:HGNC:28430] | -8.06 | 0.080 |
| *EVA1B* * | eva-1 homolog B [Source:HGNC Symbol;Acc:HGNC:25558] | -15.61 | 0.079 |
| *FAM120A* | family with sequence similarity 120A [Source:HGNC Symbol;Acc:HGNC:13247] | 7.53 | 0.051 |
| *FAM177A1* * | family with sequence similarity 177 member A1 [Source:HGNC Symbol;Acc:HGNC:19829] | -4.64 | 0.094 |
| *FAM204A* | family with sequence similarity 204 member A [Source:HGNC Symbol;Acc:HGNC:25794] | -4.78 | 0.079 |
| *GABARAPL2* | GABA type A receptor associated protein like 2 [Source:HGNC Symbol;Acc:HGNC:13291] | -4.69 | 0.051 |
| *GCFC2* | GC-rich sequence DNA-binding factor 2 [Source:HGNC Symbol;Acc:HGNC:1317] | -7.55 | 0.078 |
| *GTF2B* | general transcription factor IIB [Source:HGNC Symbol;Acc:HGNC:4648] | -3.70 | 0.095 |
| *HPSE* | heparanase [Source:HGNC Symbol;Acc:HGNC:5164] | -23.41 | 0.078 |
| *JPH2* | junctophilin 2 [Source:HGNC Symbol;Acc:HGNC:14202] | 25.59 | 0.097 |
| *JUND* * | JunD proto-oncogene, AP-1 transcription factor subunit [Source:HGNC Symbol;Acc:HGNC:6206] | -14.28 | 0.078 |

| Gene Symbol | Gene Description | Percent Change | FDR |
|---|---|---|---|
| *MAGEF1* | MAGE family member F1 [Source:HGNC Symbol;Acc:HGNC:29639] | -5.56 | 0.042 |
| *MAP1LC3A* | microtubule associated protein 1 light chain 3 alpha [Source:HGNC Symbol;Acc:HGNC:6838] | -10.25 | 0.078 |
| *MAZ* | MYC associated zinc finger protein [Source:HGNC Symbol;Acc:HGNC:6914] | 7.10 | 0.031 |
| *MICAL3* | microtubule associated monooxygenase, calponin and LIM domain containing 3 [Source:HGNC Symbol;Acc:HGNC:24694] | 17.73 | 0.078 |
| *MINK1* | misshapen like kinase 1 [Source:HGNC Symbol;Acc:HGNC:17565] | 5.01 | 0.078 |
| *MPC2* | mitochondrial pyruvate carrier 2 [Source:HGNC Symbol;Acc:HGNC:24515] | -7.73 | 0.079 |
| *MRPS14* | mitochondrial ribosomal protein S14 [Source:HGNC Symbol;Acc:HGNC:14049] | -5.61 | 0.019 |
| *MYH9* | myosin heavy chain 9 [Source:HGNC Symbol;Acc:HGNC:7579] | 7.42 | 0.034 |
| *NDUFB6* | NADH:ubiquinone oxidoreductase subunit B6 [Source:HGNC Symbol;Acc:HGNC:7701] | -5.52 | 0.094 |
| *NDUFC1* * | NADH:ubiquinone oxidoreductase subunit C1 [Source:HGNC Symbol;Acc:HGNC:7705] | -7.88 | 0.025 |
| *NFKB1* | nuclear factor kappa B subunit 1 [Source:HGNC Symbol;Acc:HGNC:7794] | 6.44 | 0.079 |
| *PKM* * | pyruvate kinase M1/2 [Source:HGNC Symbol;Acc:HGNC:9021] | 13.75 | 0.019 |
| *PRRC2B* | proline rich coiled-coil 2B [Source:HGNC Symbol;Acc:HGNC:28121] | 8.09 | 0.095 |
| *PTRHD1* | peptidyl-tRNA hydrolase domain containing 1 [Source:HGNC Symbol;Acc:HGNC:33782] | -9.26 | 0.078 |
| *RPL21* | ribosomal protein L21 [Source:HGNC Symbol;Acc:HGNC:10313] | -6.57 | 0.044 |
| *SEC11C* | SEC11 homolog C, signal peptidase complex subunit [Source:HGNC Symbol;Acc:HGNC:23400] | -8.93 | 0.031 |
| *SF1* | splicing factor 1 [Source:HGNC Symbol;Acc:HGNC:12950] | 3.72 | 0.094 |
| *SLX1B* | SLX1 homolog B, structure-specific endonuclease subunit [Source:HGNC Symbol;Acc:HGNC:28748] | -9.02 | 0.085 |
| *TAF9* | TATA-box binding protein associated factor 9 [Source:HGNC Symbol;Acc:HGNC:11542] | -4.58 | 0.100 |
| *TIMM17A* | translocase of inner mitochondrial membrane 17A [Source:HGNC Symbol;Acc:HGNC:17315] | -4.33 | 0.078 |
| *TRIM44* * | tripartite motif containing 44 [Source:HGNC Symbol;Acc:HGNC:19016] | 6.92 | 0.051 |
| *YARS1* | tyrosyl-tRNA synthetase 1 [Source:HGNC Symbol;Acc:HGNC:12840] | 4.84 | 0.078 |

* indicates a gene also identified as a DEG in the primary analysis.
